# Algostim: A breathing-synchronized neuromuscular electrical stimulation algorithm for addressing respiratory impairments after cervical spinal cord injury

**DOI:** 10.64898/2026.04.22.720073

**Authors:** Thibaut Coustillet, Nicolas Wattiez, Enora Chaïb, Adina E. Draghici, Isabelle Vivodtzev

**Affiliations:** Sorbonne Université, CNRS UMR8263, Inserm U1345, Development, Adaptation and Ageing, Dev2A, Institut de Biologie Paris-Seine, IBPS, F-75005 Paris, France; Sorbonne Université, Inserm, UMRS1158, Neurophysiologie Respiratoire Expérimentale et Clinique, F-75013 Paris, France; Cardiovascular Research Laboratory, Spaulding Hospital Cambridge, Cambridge, MA, USA; Schoen Adams Research Institute at Spaulding Rehabilitation, Boston, MA, USA; Department of Physical Medicine and Rehabilitation, Harvard Medical School, Boston, MA, USA

**Keywords:** Ordinary Differential Equation, Mathematical Modeling, Neuromuscular Electrical Stimulation, Spinal Cord Injury, Breathing Dysfunction

## Abstract

Cervical spinal cord injuries (cSCI) induce profound denervation in respiratory muscles leading to hypoventilation that compromises quality of life. Respiratory neuromuscular electrical stimulation of extra-diaphragmatic muscles (rNMES) could be a non-invasive approach to improve respiratory function following cSCI. However, asynchronous NMES with spontaneous respiration can feel unnatural or painful or even lead to potential complication, although synchronized stimulation may be more efficient to improve neuroplasticity after SCI. Here we developed a software-driven synchronized rNMES system aligned with spontaneous breathing, with preliminary validation in a mouse model of cSCI.

**Methods:** An Ordinary Differential Equation (ODE) was solved and fitted to experimental breathing signals obtained via respiratory function recording in mice which underwent cSCI. Optimal stimulation ODE-based parameters were then identified for both intercostal and abdominal muscle stimulation for breathing-synchronized rNMES training. Lastly, acute efficacy was assessed by evaluating the increase in chest position during intercostal stimulation, measured using a piezoelectric sensor on the thorax during stimulation.

**Results:** The ODE-based breathing signals matched the experimental ones with an average coefficient of determination (R²) of 81%. The developed algorithm, *Algostim*, provided average theoretical optimal times of 0.10 ± 0.01 s for intercostal and 0.17 ± 0.02 s for abdominal muscles contraction. Synchronized intercostal stimulation led to twice greater change in chest amplitude compared to non-synchronized stimulation and there was a dose dependent effect with a minimal current intensity of 2 mA and an optimal stimulation reached with 3 mA. This innovative mathematical approach can optimize respiratory muscle stimulation while accounting for spontaneous breathing rate, establishing a framework for personalized rNMES therapies and enabling investigation into its underlying mechanisms.

## 1 Introduction

The majority of spinal cord injuries (SCI) occur at the cervical level, resulting in marked respiratory dysfunction^1^. Disruption of spinal cord integrity leads to profound reductions in descending supraspinal respiratory pathways, with the severity usually proportional to the level and extent of injury. A complete lesion at or above the level of the 4^th^ cervical vertebra induces phrenic motoneurons alteration and/or denervation resulting in diaphragmatic paralysis and subsequent loss of spontaneous breathing^2^. In addition, the impact of cervical SCI (cSCI) extends to intercostal and abdominal muscles that cannot compensate for reduced diaphragmatic contraction^3^.

Currently there is no treatment to restore respiratory function in cSCI. Almost all tetraplegic individuals require mechanical ventilation acutely after injury, with many remaining dependent on it thereafter. Even when those with cSCI are weaned from diurnal mechanical ventilation, most (60%) still require nocturnal ventilation support due to sleep disordered breathing^4^. Implanted phrenic stimulation (diaphragm pacing) allows some individuals to be weaned from mechanical ventilation^5^. However, this invasive approach is only possible when phrenic nerve conduction is preserved, which occurs in less than 10% of SCI with lesions above C4^6^. Thus, there is an urgent need to develop alternative interventions to increase the number of adults with cSCI eligible for phrenic nerve implantation.

Neuromuscular electrical stimulation (NMES) is a technique employing intermittent electrical stimuli to induce muscle contractions *via* surface electrodes placed on the skin. NMES has been successfully employed for over a decade to mitigate muscle wasting^7,8^. In addition, NMES targeting extra-diaphragmatic respiratory muscles (rNMES), such as intercostal and abdominal muscles, has been shown to increase respiratory function after cervical SCI^9,10^. When applied to the pectoralis major and abdominal muscles, rNMES results in increased chest amplitude and cough capacity in adults with SCI. However, despite these promising results, the molecular mechanisms underlying the observed improvements following (r)NMES interventions remain largely unexplored. Some pathways might be involved in enhancing neuroplasticity, potentially facilitating spontaneous recovery of phrenic nerves. Of note, NMES favors the recovery of mechano- and metabo-sensitivity of denervated muscle in rats^11^. Moreover, similar to exercise, NMES increases plasma levels of brain-derived neurotrophic factor^12,13^, well-known to promote neurogenesis, neuroplasticity, and synaptogenesis^14^. Hence, NMES may be a promising experimental approach for neuromuscular regeneration^15^ as well, particularly when applied to respiratory muscles in individuals with cSCI reliant on mechanical ventilation. However, its efficiency must be validated through pre-clinical studies to confirm neuroplasticity before clinical applications in people dependent on ventilatory support. A significant challenge of current rNMES systems is their limited ability to adapt to rapid breathing rates of rodents, which can reach up to 200 breaths per minute. Furthermore, to date, the parameters for rNMES stimulation, including contraction and rest times, are largely empirical and do not account for intrinsic breathing pattern: there is a lack of consensus and reproducibility.

Thus, overcoming these limitations and developing an optimized pre-clinical rNMES stimulation approach well-adapted to breathing dynamics is a crucial first step before evaluating its efficiency as a potential therapy for SCI rehabilitation. Hence, we sought to develop a rNMES approach that integrates the breathing patterns of mice following cervical SCI, ensuring optimal synchronization between rNMES and spontaneous breathing. We previously developed a mouse model of cervical hemi-contusion to serve as a pre-clinical model of cervical SCI for interventional studies^16^. Additionally, we previously established a chronic, *in vivo* model of NMES of the lower limb in mice^17^.

Here, we developed *Algostim*, an algorithm that employs an ordinary differential equation (ODE) to model breathing. Afterwards, in order to determine individualized stimulation parameters for each animal, the model was fitted with experimental breathing patterns obtained *via* plethysmography. Moreover, we demonstrated the acute efficacy of synchronized intercostal rNMES by evaluating the effect of synchronization with respiratory cycles on chest expansion, as well as the dose-response relationship of increasing current intensity during intercostal stimulation. This was measured using a piezoelectric sensor on the thorax of mice after cervical hemi-contusion. To our knowledge, this represents the first analytical approach of breathing integrating experimental physiological signals in mice following cervical hemi-contusion or sham surgery.

## 2 Methods

### 2.1 Experimental data

#### 2.1.1 Animals

Experiments were made on (8-10-week-old, 36 ± 3 g) male wild-type mice (*Mus musculus*, swiss OF-1 strain; Charles River laboratories, France). Animals were housed in cage of five and were accustomed to live together one week before the beginning of the study and in cage of two after surgery for better control of post-surgery care. Alimentation and environment (space, light and noise) was strictly similar between cages. Animals were kept on a 12 h light-dark cycle with free access to food and water. The study was conducted in accordance with the Directive 2010/63/EU of the European Parliament and of the Council of 22 September 2010 and French law (2013/118), and to the Guide for Care and Use of Laboratory Animals (NIH Publication No. 85–23, revised 1996). The protocol was approved by the French ethical committee CEEA – 005, Charles Darwin, with ethical approval statement #26668 – 2020070317412138 v3, November 2020). All efforts were made to minimize suffering and number of animals used.

#### 2.1.2 cervical Hemi-Contusion (cHC)

Hemi contusion models have been performed as previously described in our previous work^16^. Briefly, animals were anesthetized with isoflurane (1.5 – 2.5 %) throughout the procedure *via* a facial mask. Dorsal skin and underlying muscles from the second cervical to the first thoracic vertebrae were retracted. A dorsal laminectomy was performed to expose the spinal cord. A manual impactor device was used with a 1.0 mm tip impactor (RWD life science; 68,099 II). After contusion, sutures were performed to close the wounds and skin and post-surgery care were applied to the animals.

#### 2.1.3 Breathing measurements

Spontaneous breathing was obtained *via* non-invasive whole-body plethysmography as previously described^16^. Briefly, awake animals were placed in 250 mL chambers, breathing spontaneously and unconstrained (Emka, France). Success rate (Sr) was calculated by the acquisition software as the ratio of the number of validated cycles to the number of detected cycles for each period of 10 s. A Sr = 100% indicates that all detected respiratory cycles are validated with the highest confidence by the plethysmography software (IOX software, Emka, France). Thus, the criteria for inclusion in mathematical modeling consisted of a minimum Sr of 60% as well as a manual curation of each plethysmography recording to ensure adequate stability of the recorded signal and relevant stimulation parameters.

### 2.2 Study design

The study design is presented in Figure 1. In this study, 20 mice (cHC, n = 16; laminectomy, n = 4) were acclimatized for four days to the plethysmography chambers (d-7, d-3) for one hour a day to familiarize themselves with chambers. Spontaneous breathing (raw air flow rate) was recorded *via* plethysmography three days before surgery after proper acclimatization of the animals (d-3). After 7 days of rest and care following surgery (d0), a second recording of spontaneous breathing was obtained *via* plethysmography (d7). Out of the initial cohort of 20 mice, n = 10 (n = 8 with cHC and n = 2 with laminectomy) had plethysmography signals of adequate quality both before and after surgery (Quality score ≥ 65%), ensuring appropriate mathematically modeling of their breathing patterns. Among the remaining 10 mice, n = 2 died due to complications after cHC, and n = 8 mice did not reach the signal quality requested for mathematical modeling but were used to test the acute effect of rNMES, based on the calculation of optimal rNMES parameters using *Algostim*.

**Figure 1:**
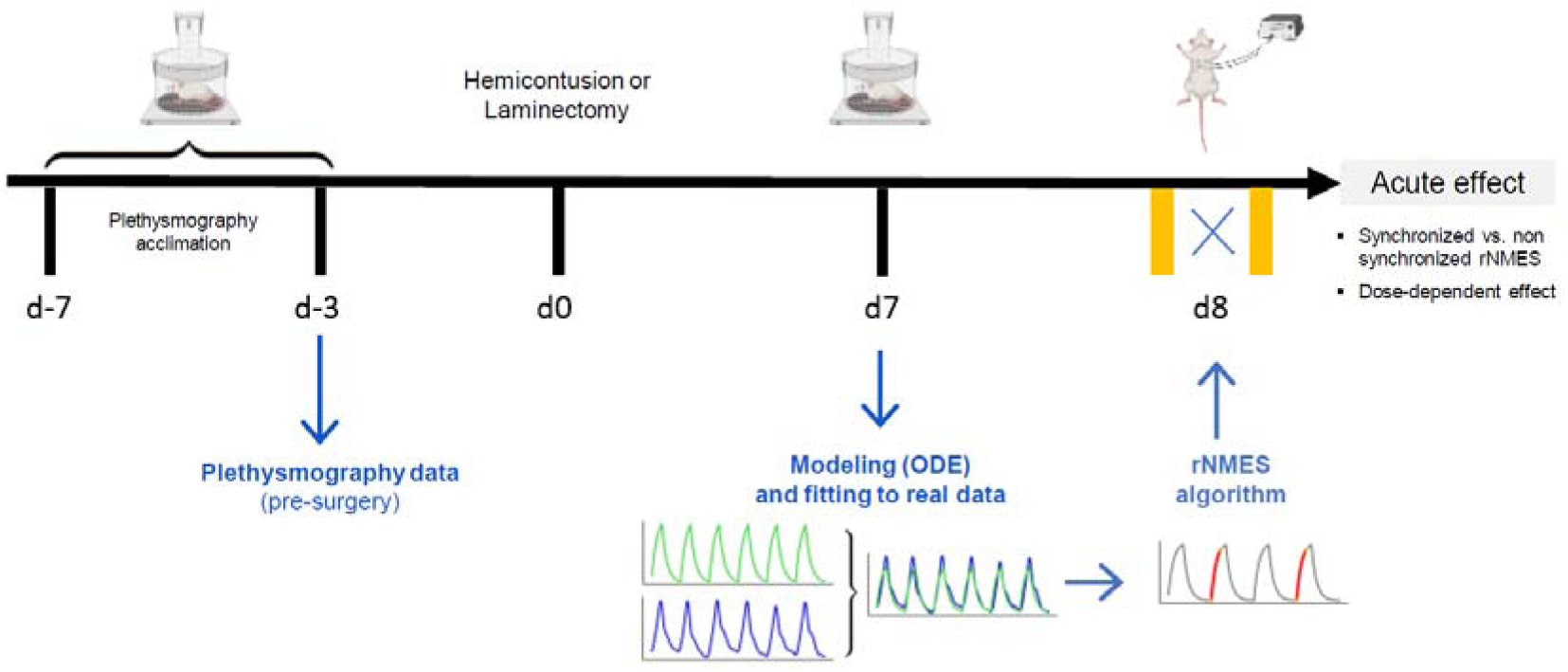
Study design. Milestones from acquisition of experimental breathing signals to determination of individualized rNMES stimulation parameters and rNMES testing sessions. cHC, cervical hemi-contusion, ODE, Ordinary Differential Equation; rNMES, respiratory neuromuscular electrical stimulation.

### 2.3 Mathematical model of breathing

A theoretical breathing signal was derived employing a mathematical model of breathing predicated on an Ordinary Differential Equation (ODE). The optimal ODE solution (*i.e.*, theoretical breathing) was obtained by a least-squares fit with the experimental breathing signals obtained *via* plethysmography. Subsequently, the adjusted ODE breathing model was used to determine optimal parameters for breathing-synchronized NMES. Throughout the study, we refer to (1) the first part as the ODE solution and (2) the optimal theoretical parameter calculation part as the algorithm (*Algostim*). ODE-based breathing signal being the fitting of the ODE solution to experimental data, it was later referred to as the theoretical breathing.

#### 2.3.1 The pipe-balloon model

Based on Maury’s previous work, the respiratory tract was considered as a single pipe (no sub-branches) of constant resistance R (*i.e.*, R ⋲ □) connecting alveoli to ambient air, where all the alveoli were represented as a unique balloon^18^. This simplified model, termed as the *pipe-balloon*, ensures communication between the elastic balloon (representing the lungs) and the external environment (Fig. 2).

**Figure 2:**
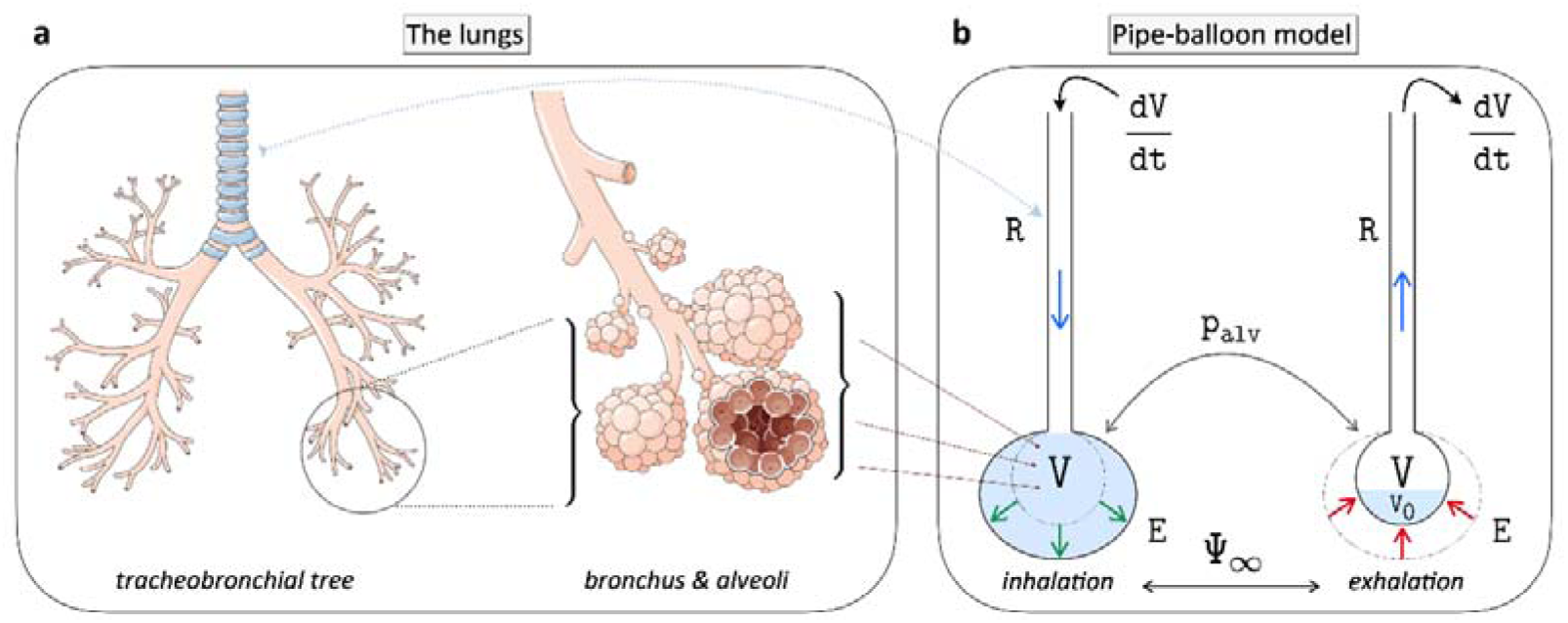
The pipe-balloon model. (a) Simplified representation of the lungs. (b) Comprehensive model used in the study. The Ordinary Differential Equation derives directly from the pipe-balloon model. dV/dt, Air flow rate; E, Elastance; p_alv_, alveolar pressure (balloon pressure); R, Resistance; V, Volume; V_0_, Residual volume; □_∞_, Periodicity of respiratory muscles. Panel (a) was designed from elements supplied by Servier Medical Art (https://smart.servier.com/), licensed under a Creative Commons Attribution 4.0 Unported License.

Periodicity of respiratory muscles. Panel (a) was designed from elements supplied by Servier Medical Art (https://smart.servier.com/), licensed under a Creative Commons Attribution 4.0 Unported License.

The pipe-balloon model (Fig. 2b) is based on Poiseuille’s law stating that the flow velocity of a fluid between two points is proportional to pressure difference between them. Thus, in the pipe-balloon model, the airflow rate dV/dt, is proportional to the pressure difference between the balloon (≔ p_balloon_) and the ambient air (≔ p_atm_). Considering p_balloon_ = p_atm_ + p_alv_, where p_alv_ is the internal pressure of the balloon, the pressure difference is given by the following equation:

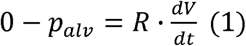

At rest, the capacity of the lungs to fill with air depends on the pressure exerted by inspiratory muscles, mainly the diaphragm, but also on extra-diaphragmatic muscles such as (external) intercostal muscles. Indeed, inspiratory muscle contraction leads to rib cage expansion, increased chest volume, and subsequently to a drop-in pressure in the pulmonary alveoli^19^. As alveolar pressure is lower than atmospheric pressure, the air comes inward following its natural pressure gradient. Hence, the more the inspiratory muscles contract, the more the lungs can fill with air. In addition, considering the histological and physiological properties of the lungs, their elasticity is a function of elastance E (assumed linear relationship) and residual volume after active exhalation (V_0_). Thus, the deformation behavior of the balloon is given by:

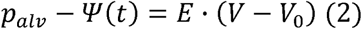

Where □ denotes the pressure induced by inspiratory muscles contraction and V denotes the balloon volume at all times t ⋲ □_+_. By adding equations (1) and (2), we obtained the following ODE:

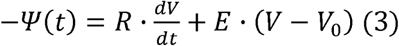

Assuming that resting exhalation was passive (only through the relaxation of inspiratory muscles), the analytical expression of the function □ (equation 4) during a single breathing cycle (of duration denoted T_cycle_) only depends on the negative pressure □⋲ □_-_ exerted during inhalation (depression, from 0 to inspiratory time = T_insp_). There is no activity during the exhalation (*i.e.*, □ α 0 from T_insp_ to T_cycle_).

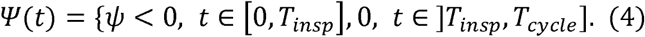

□ was simply determined as the average amplitude (tidal volume; V_T_) of a given measured experimental signal (≔ A(s)) derived from plethysmography. We assumed that increased amplitude of the breathing signal is directly correlated with greater inspiratory muscle contraction, resulting in greater magnitude of in absolute value.

Given that the pressure exerted by the inspiratory muscles is negative (depression), □ for each mouse is defined as:

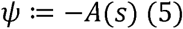

Where A is a function that returns the average amplitude of an experimental signal s. Similarly, T_insp_ and T_cycle_ are deducted as the inspiratory time and the duration of one complete breath (one inhalation + one exhalation) of the above-mentioned s signal. As we modelled on 10 mice, there were 10 different functions □ (10 signals).

Lastly, the periodic activity of the inspiratory muscles, □_∞_, represented all the periodic breathing cycles induced by the respiratory muscles with a period T_cycle_. Specifically, □_∞_ was defined as a periodic function of period T_cycle_ given by:

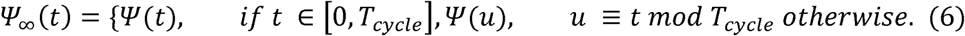

Considering the periodic contractions of the inspiratory muscles and rearranging equation (3), the final linear ODE is obtained as shown in equation 7. The ODE solving represents the first part of *Algostim* allowing subsequent derivation of optimal parameters to inform rNMES stimulation (second part) in the present study.

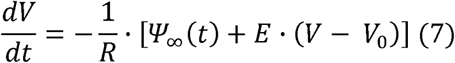

where R, E, and V_0_ represent constant values of elastance, resistance, and residual volume, respectively.

#### 2.3.2 Comparison of theoretical and experimental breathing signals (fitting)

For each mouse (n=10), experimental breathing signals obtained *via* plethysmography (Fig. 1) were used to derive the ODE solutions (theoretical signals). Breathing signals exported from Emka technologies software were digitized at 500 Hz to ensure good accuracy with reasonable computing time. Afterwards, Savitzky-Golay (SG) filter was used to reduce any spontaneous breathing noise. Briefly, the SG filter is based on subdivision of signals in regular time intervals (length of 47 points here) and, for each interval, a polynomial (of degree 3) regression was used to minimize errors based on least-squares fit^20^. Given that Emka software provided only instantaneous flow (dV/dt), the volume was computed by integrating the filtered breathing signals using the trapezoid method^21^.

The ODE was then solved using the *odeint* solver in python 3.X. Similarly to the SG filtering, the constant values (R, E and V_0_) were determined by minimizing errors *via* least-squares method between experimental breathing signals and ODE solutions. Specifically, the optimal solution of equation (7), also denoted as the theoretical breathing signal, depends on the function V(t, ξ_min_) with ξ_min_ = (R_min_, E_min,_ V_0min_) ɛ R^3^given by:

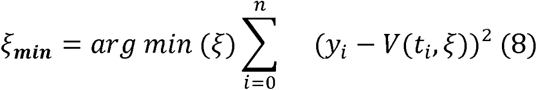

where y_i_ denotes the i^th^ experimental point of a given experimental signal and V(t_i_, ξ) the i^th^ theoretical derived value given the ξ solution.

#### 2.3.3 Refinement of theoretical breathing accounting for the effect of anesthesia

Given that stimulation training is conducted while on anesthetized animals and considering the known effects of anesthesia on dampening breathing frequency^22^ (ƒ_B_), it was essential to account for the frequency of the theoretical breathing signal prior to determining the optimal durations for breathing-synchronized rNMES. Thus, the effect of anesthesia was measured on eight animals not included in the initial modeling. Data obtained at 5 min and at 30 min post anesthesia were used to determine the ƒ_B_ characteristics over 30 min of anesthesia. Thus, the effect of anesthesia on ƒ_B_ was given by a dilation factor calculated as the ratio of the initial ƒ_B_ (prior to anesthesia) and the ƒ_B_ at time t after t min of anesthesia as follows:

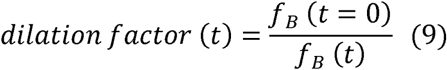

where t ⋲ [0, 30] the time of anesthesia.

#### 2.3.4 Determination of optimal contraction and rest times for rNMES training

From the refined theoretical breathing signals accounting for the effect of anesthesia, optimal contraction and rest durations were computed to ensure that the stimulation time did not exceed 25% of the total [contraction + rest] cycle. Additionally, the algorithm was designed to allow a fix number of unstimulated breaths between stimulation cycles which can be freely chosen. For each mouse, tetanic contraction times were set at 65% of the inspiratory phase for intercostal muscles and 75% of the expiratory phase for abdominal muscles. This approach guarantees synchronization with spontaneous breathing, regardless of phase-specific stimulation ratios or the chosen number of rest cycles, while preserving full inhalation and exhalation dynamics^23^.

### 2.4 Acute effect of rNMES

Height animals underwent intercostal rNMES training sessions to test its ability to increase chest amplitude. Two parameters were tested: the effect of synchronization vs. non-synchronized stimulation and the effect of current intensity level. A cross-over design with randomized order of first occurrence was used to test the effect of synchronized vs. non-synchronized stimulation and for synchronized stimulation, current intensity was then progressively increased from 0 to 4 mA every 1 mA. Animals were anesthetized using Isoflurane inhalation at a concentration of 2% and lying on a heating pad to maintain normal core body temperature. Animals underwent skin depilation and a conductive universal gel was applied between the electrodes and the skin to facilitate effective current conduction. Electrical stimulation was delivered using a symmetrical, biphasic, square-pulsed electrical current with a voltage of 15 V for the contraction periods and 0 V for the rest periods, with a tetanic pulse width duration of 8 ms (digitimer, model DS7A; PowerLab®, Labchart pro 8.1.16, ADInstruments). The contraction and rest time durations for each mouse were based on the stimulation parameters derived with *Algostim*. rNMES training testing was performed7 days after surgery (Fig. 1). Measurements during stimulation were performed using a piezo electrode positioned on the thorax rostrally to the stimulation electrodes which were transduced and integrated on the PowerLab®, Labchart pro 8.1.16, ADInstruments). The piezo sensor detects thoracic movements induced by breathing, converting mechanical deformations into an electrical signal proportional to the chest’s expansion and contraction. In one mouse, the piezo sensor signal was too poor to obtain reliable data preventing us from accurately assessing the effect of stimulation.

## 3 Results

### 3.1 ODE solution and validation (fitting)

Assuming an initial volume V_0_ at time t=0, the exact solution of equation (7) is given by^18^:

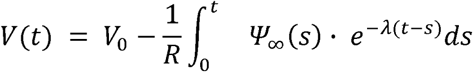

with λ=E/R.

The ODE-based breathing signal represents the theoretical breathing an animal would have if breathing always remained constant, i.e., if each breathing cycle was strictly identical to its neighbor. Figure 3 outlines a part of the framework (without the refinement, which covers the anesthesia) from experimental signal acquisition to fitting and calibration of the theoretical signal (1^st^ part of *Algostim*). Following experimental data acquisition (Fig. 3a), the raw air flow rate (dV/dt) signal obtained *via* plethysmography (Fig. 3a) was denoised *via* SG filtering (Fig. 3b) and the volume V was calculated by integrating the smoothed air flow rate over time (Fig. 3c). Afterwards, the theoretical signal (ODE solution) (Fig. 3d) was fitted to the experimental breathing signal *via* least-squares method (Fig. 3e).

**Figure 3:**
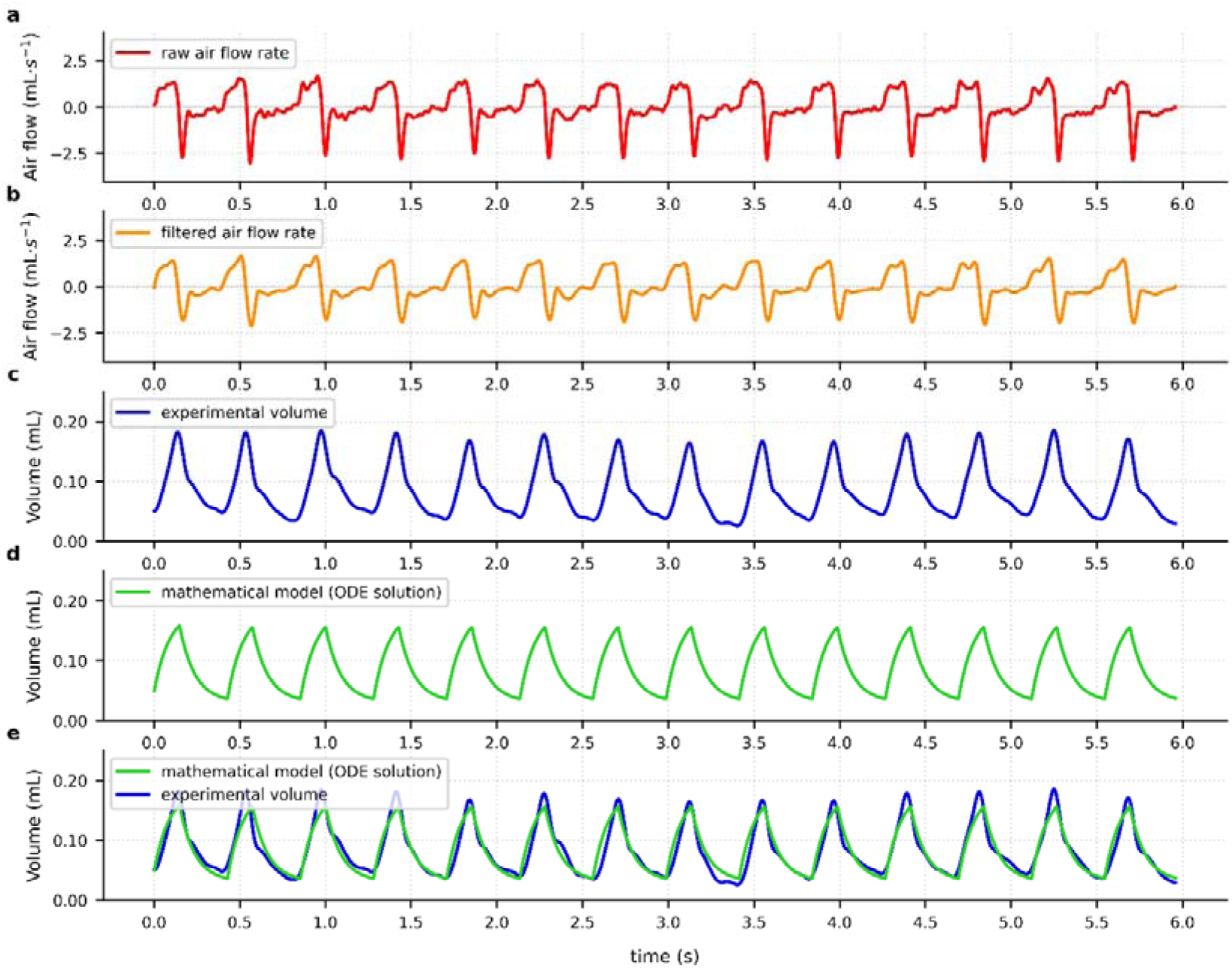
Framework for deriving theoretical breathing signal through ODE solution incorporating experimental plethysmography data. The five panes refer to the framework for a single animal. (a) raw air flow rate, (b) filtered air flow rate, (c) integrated air flow rate, (d) theoretical volume, (e) merge between integrated air flow rate and theoretical volume.

Individual theoretical breathing signals were derived for each of the 10 mice. On average, the theoretical breathing signals corresponded closely to the experimentally obtained breathing signals (Fig. 3e), with a coefficient of determination (R^2^) of 81 ± 5 %. Similarly, on average, theoretical ƒ_B_ derived from the theoretical breathing signals was similar to the experimental one (Table 1; 156 ± 24 rpm vs. 155 ± 24 rpm). However, there were more noticeable differences between theoretical and experimental V_T_ amplitudes (Table 1; 0.14 ± 0.02 mL vs. 0.17 ± 0.02 mL). The mean approximation error was 0.5 ± 0.4 % for ƒ_B_ and 14.2 ± 9.9 % for V_T_ amplitude, resulting in a total approximation error of 7.4 % when considering these two parameters alone. The discrepancy in amplitudes can be attributed to the inherent constraints of the ODE model, including the linear relationship between balloon volume and elastance, which assumes a concave waveform for inhalation and a convex waveform for exhalation. This assumption prevents an exact fit of the peak volume between theoretical breathing signals and experimental data (Fig. 3d-e). Nonetheless, the impact of this amplitude difference is negligible given that the algorithm for determining optimal parameters relies solely on ƒ_B_ (inspiratory and expiratory times) and not on signal amplitude. Lastly, analysis of the 10 theoretical breathing signals resulted in a mean resistance value of R = 39 ± 3 (no unit), a mean elastance value of E = 0.7 ± 0.3 (no unit) and a mean residual volume value of V_0_ = 0.04 ± 0.03 mL (Table 1).

**Table 1:** Overview of respiratory features for ODE solutions. Goodness-of-fit given by R^2^ and approximation errors. Approximation errors were calculated as 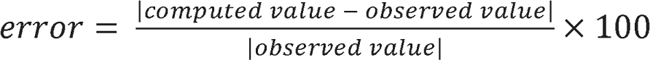. cHC, cervical hemi-contusion; E, Elastance; ƒ_B_, frequency of breathing; LM, laminectomy; R, Resistance; rpm, respiration per minute; Sr, Quality score; V_T_, Tidal volume; V_0_, Residual volume.

| Mouse | Group | Sr of experimental signal (%) | Experimental $f_B$ (rpm) | Theoretical $f_B$ (rpm) | $f_B$ approximation error (%) | Experimental $V_T$ amplitude (mL) | Theoretical $V_T$ amplitude (mL) | $V_T$ amplitude approximation error (%) | Overall error (%) | $R^2$ (%) | R | E | $V_0$ |
| --- | --- | --- | --- | --- | --- | --- | --- | --- | --- | --- | --- | --- | --- |
| 1 | LM | 68 | 184 | 185 | 0.5 | 0.16 | 0.16 | 0.0 | 0.3 | 81 | 36 | 0.3 | 0.02 |
| 2 | LM | 100 | 141 | 141 | 0.0 | 0.14 | 0.12 | 14.3 | 7.1 | 90 | 40 | 0.9 | 0.03 |
| 3 | cHC | 100 | 184 | 185 | 0.5 | 0.17 | 0.13 | 23.5 | 12.0 | 72 | 35 | 0.9 | 0.05 |
| 4 | cHC | 96 | 166 | 167 | 0.6 | 0.16 | 0.14 | 12.5 | 6.5 | 81 | 42 | 0.3 | 0.01 |
| 5 | cHC | 94 | 131 | 133 | 1.5 | 0.16 | 0.11 | 31.2 | 16.4 | 74 | 35 | 1.3 | 0.07 |
| 6 | cHC | 100 | 169 | 169 | 0.0 | 0.19 | 0.15 | 21.1 | 10.5 | 78 | 35 | 0.9 | 0.05 |
| 7 | cHC | 65 | 105 | 106 | 0.9 | 0.22 | 0.17 | 22.7 | 11.8 | 79 | 44 | 0.9 | 0.10 |
| 8 | cHC | 100 | 175 | 176 | 0.6 | 0.15 | 0.15 | 0.0 | 0.3 | 84 | 38 | 0.4 | 0.02 |
| 9 | cHC | 100 | 149 | 149 | 0.0 | 0.15 | 0.14 | 6.7 | 3.3 | 88 | 40 | 0.7 | 0.02 |
| 10 | cHC | 100 | 148 | 149 | 0.7 | 0.19 | 0.17 | 10.5 | 5.6 | 84 | 43 | 0.7 | 0.02 |
| Mean |  | 92 | 155 | 156 | 0.5 | 0.17 | 0.14 | 14.2 | 7.4 | 81 | 39 | 0.7 | 0.04 |
| Sd |  | 13 | 24 | 24 | 0.4 | 0.02 | 0.02 | 9.9 | 5.0 | 5 | 3 | 0.3 | 0.03 |

### 3.2 Refinement of theoretical breathing signal accounting for the effect of anesthesia

Anesthesia is well known to reduce breathing frequency. Thus, in the current study we accounted for the effect of anesthesia on ƒ_B_ as shown in Fig. 4. On average (n=8 anesthetized mice), ƒ_B_ decreased from 5 min (147 ± 12 rpm) to 30 min (83 ± 6 rpm). The estimated ƒ_B_ prior to anesthesia was 160 rpm. Assuming a linear decline in ƒ_B_, the dilation factor (−) increases inversely proportional with time from ∼1.1 at t=5 min and reaches a maximum of 1.94 after 30 min of anesthesia. (Fig 4a). Thus, in the present study, due to experimental challenges associated with manually updating stimulation parameters over time, we considered a constant factor of 1.5, which represents an intermediate value of the observed range during anesthesia (Fig 4b-c). Given that the main objective of this study is to demonstrate the feasibility of *Algostim* during rNMES, the constant dilation factor provides an appropriate estimate of the effect of anesthesia.

**Figure 4:**
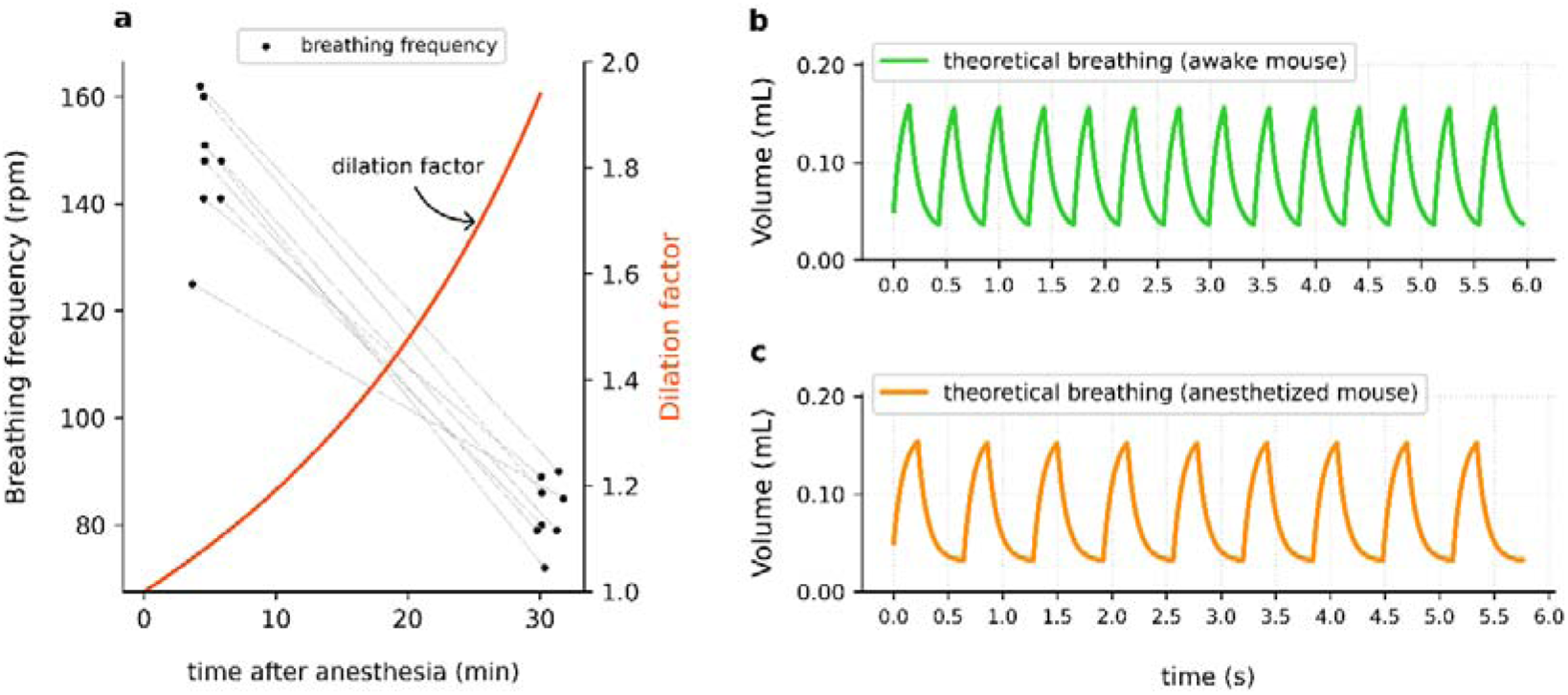
Refinement of theoretical breathing signal to account for reduced breathing frequency during anesthesia. (a) Breathing frequency of eight anesthetized mice (2% isoflurane) at two time points (5 min v 30 min). The dilation factor (orange) denotes the effect of the anesthesia. (b) Theoretical breathing signal in an awake mouse. (c) Theoretical breathing signal of the same mouse under anesthesia considering a constant dilation factor of 1.5. Both signals have the same amplitude (tidal volume).

### 3.3 Determination of optimal contraction and rest times for rNMES training (2^nd^ part of *Algostim*)

The *Algostim* framework from experimental signal acquisition to optimized parameter calculation is shown in Fig. 5. The algorithm starts with a discretized plethysmography breathing signal. The raw signal undergoes filtering and integration, followed by manual validation before advancing to subsequent steps. Failure to advance to the next step, for instance, due to poor signal quality, an alternative more suitable time window may be considered for modeling. After modeling and refinement are completed (1^st^ part), another manual checkpoint is necessary before the algorithm can determine the optimal parameters (2^nd^ part).

**Figure 5:**
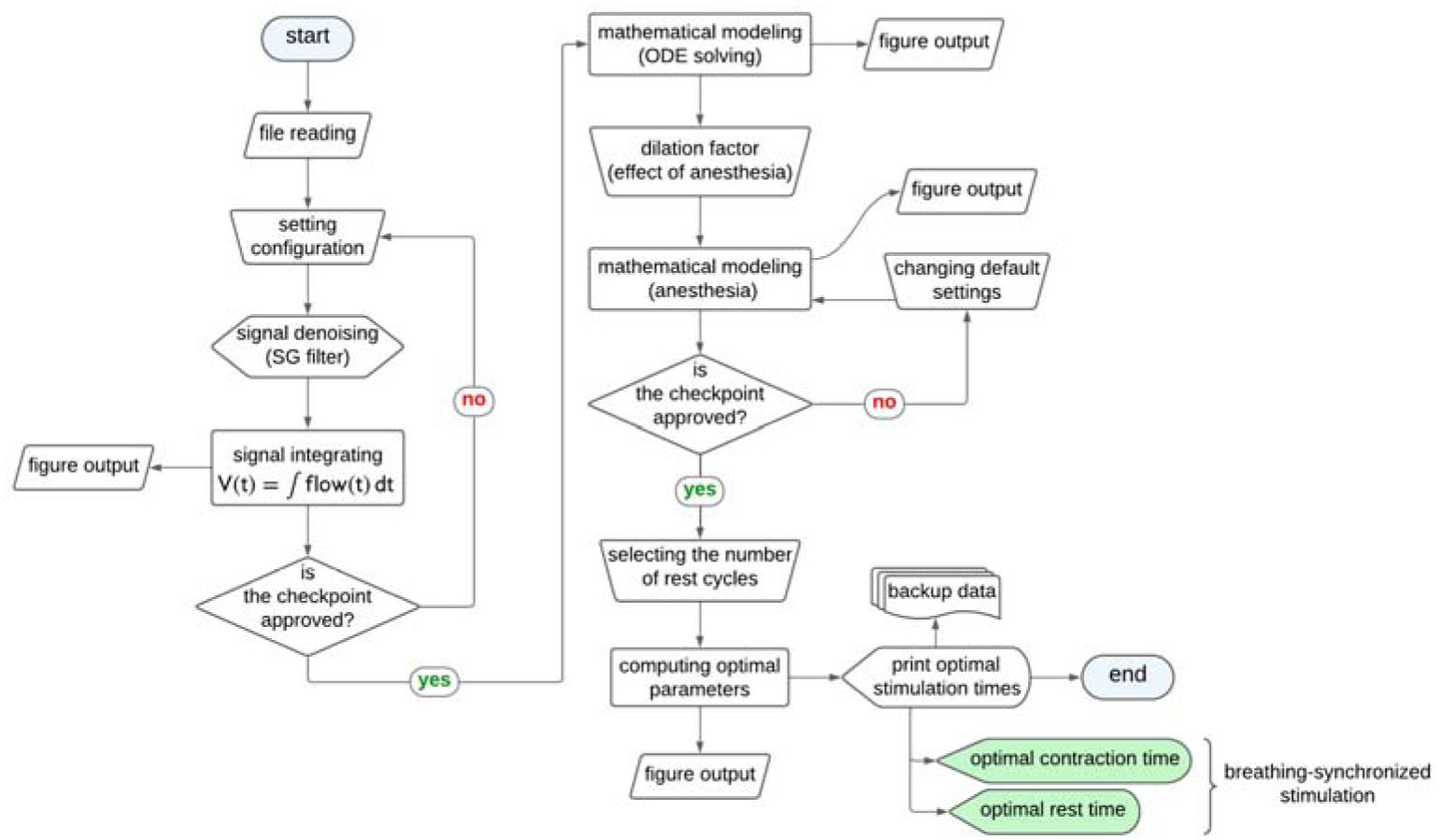
Flow chart of *Algostim* for determining optimal parameters for breathing-synchronized stimulation. Figure created in Lucid (<u>lucid.co</u>).

Individualized rNMES parameters for the stimulated mice are reported in Table 2. For intercostal muscle contractions (inhalation phase), average rNMES duration was 0.10 ± 0.02 s with 1.80 ± 0.17 s rest periods and for abdominal muscle (exhalation phase) rNMES duration was 0.17 ± 0.02 s with 1.35 ± 0.13 s rest periods Figure 6 represents breathing-synchronized theoretical stimulation for intercostal and abdominal muscles. During stimulation training, as for example, we allowed four cycles of rest between each intercostal contraction (inhalation; Fig. 6a) and three complete cycles of rest between each abdominal contraction (exhalation; Fig. 6b). Theoretical stimulation of the intercostal (external) muscles occurred only during inhalation (Fig. 6a), while theoretical stimulation of the abdominal muscles occurred only during the exhalation phase (Fig. 6b). As mentioned in the Methods section, the stimulated portions (red) accounted for 65% of the inspiratory phase and 75% of the expiratory phase.

**Figure 6:**
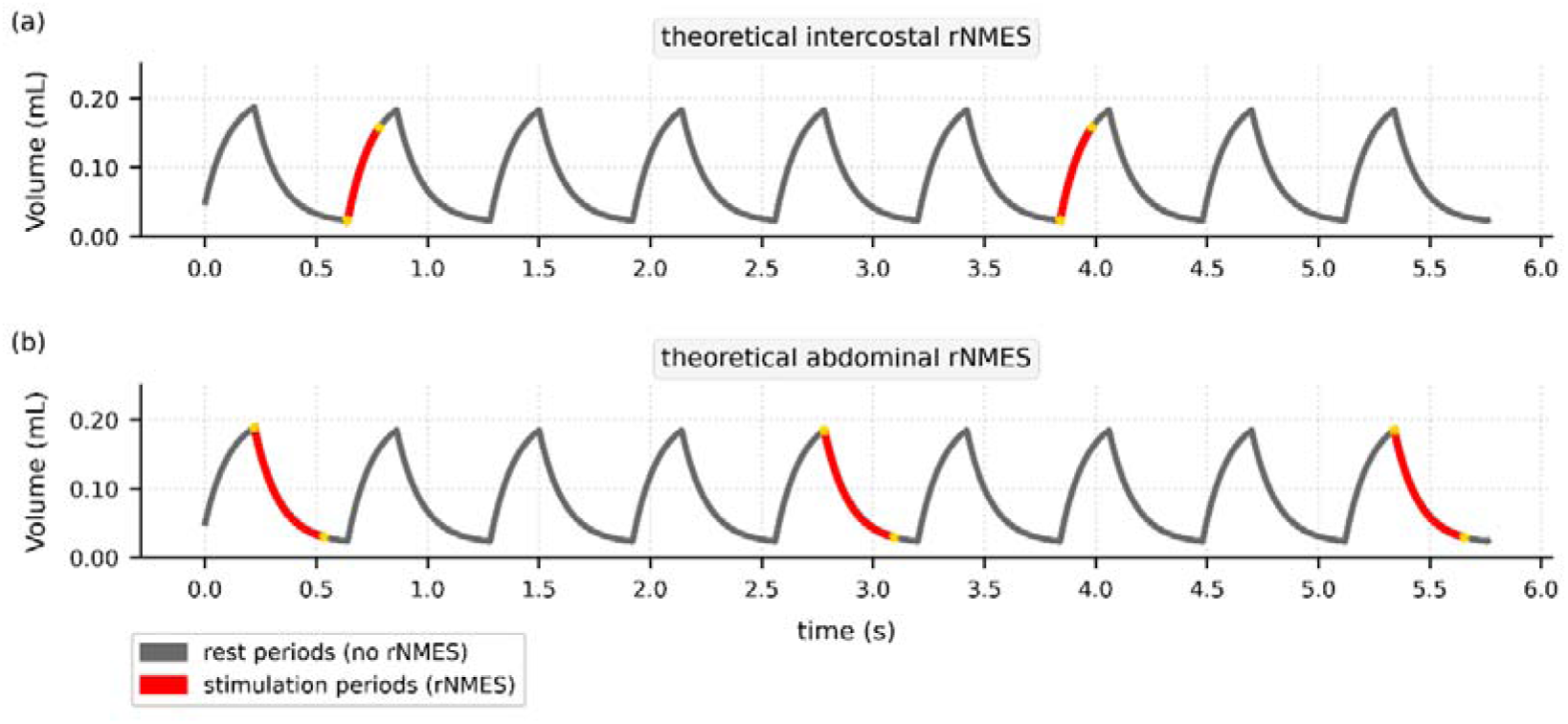
Output of the *Algostim* algorithm and theoretical contraction periods during the breathing-synchronized rNMES. Stimulation periods are depicted in red and rest periods are shown in gray. Although 6 s are presented for convenience, the stimulation lasted 15 min for each muscle group.

**Table 2:** Overview of respiratory parameters for Synchronized rNMES (Algostim) used for acute testing in cHC mice.

|  | Intercostal muscles |  | Abdominal muscles |  |
| --- | --- | --- | --- | --- |
| Mouse | Contraction time | Rest time | Contraction time | Rest time |
| 11 - cHC | 0.10 | 1.88 | 0.18 | 1.4 |
| 12 - cHC | 0.08 | 1.53 | 0.15 | 1.14 |
| 13 - cHC | 0.11 | 1.95 | 0.19 | 1.46 |
| 14 - cHC | 0.11 | 1.98 | 0.19 | 1.48 |
| 15 - cHC | 0.12 | 1.76 | 0.15 | 1.36 |
| 16 - cHC | 0.09 | 1.62 | 0.16 | 1.21 |
| 17 - cHC | 0.10 | 1.88 | 0.18 | 1.4 |
| **Mean** | **0.10** | **1.80** | **0.17** | **1.35** |
| Sd | 0.01 | 0.17 | 0.02 | 0.13 |

### 3.4 Acute effect of synchronized rNMES (***Algostim***)

Figure 7 shows acute response to rNMES as assessed by change in chest position over time (figure 7A). When well synchronized with spontaneous breathing, stimulation led to an increase in chest position significantly higher than without stimulation or non-synchronized stimulation and almost twice higher than non-synchronized stimulation (p = 0.001, Figure 7B). Moreover, we found a dose-effect relationship with a minimal current intensity of 2 mA to induce a significant increase in chest position and an optimal current intensity (efficient and tolerable) of 3 mA for which change in chest position started to plateau or current started to be intolerable. Of note, on mouse did not tolerated current intensity ≥ 3 mA and one did not tolerate current intensity ≥ 4 mA.

**Figure 7:**
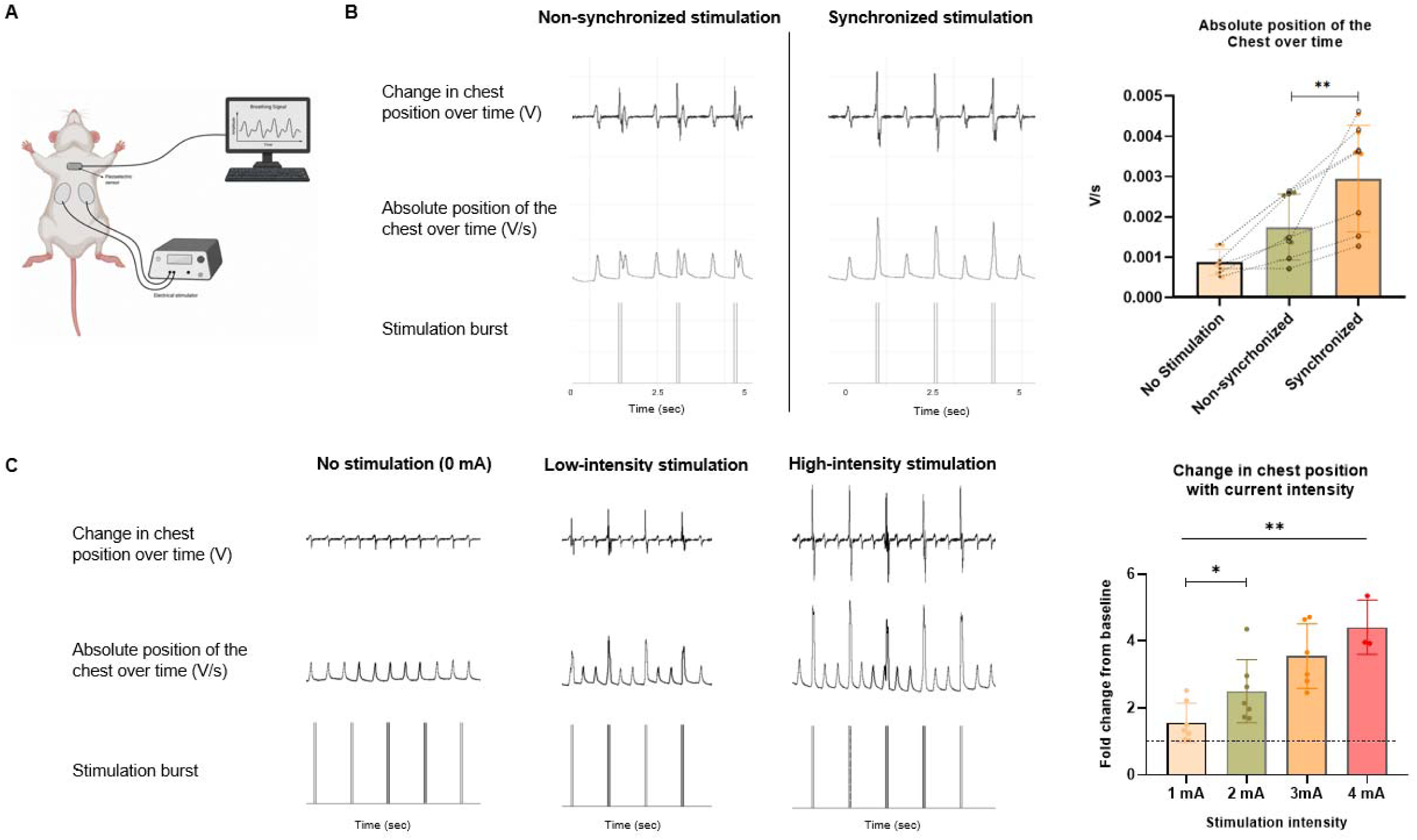
Acute effect of rNMES. A/ Schematic representation of the experimental setting to evaluate chest position. B/ Change in chest position over time while intercostal stimulation is synchronized or not to spontaneous breathing. C/ Effect of current intensity level on change in chest position over time with synchronized intercostal stimulation. Data is presented as mean ± sd.

## 4 Discussion

Here, we developed a mathematical model of breathing and used experimental data to establish an algorithm (*Algostim)* designed to facilitate breathing-synchronized stimulation in a mouse model of cervical SCI. *Algostim* is based on the ‘*pipe-balloon*’ model and mimics breathing by leveraging experimental data. We have shown that *Algostim* determines optimal rNMES parameters including duration and timing of intercostal and abdominal muscle contraction during spontaneous breathing in mice. This approach provides individualized stimulation parameters aimed at increasing the structure and function of respiratory muscles without inducing inconsistent respiratory movements during spontaneous breathing. The framework is designed to deliver a method for calculating stimulation parameters that is both objective and reproducible, thereby avoiding the use of arbitrary “round” values that lack clear physiological relevance (*e.g.*, exactly 1 s or 1.5 s).

Current mathematical models describing respiratory function are based on ODE or Navier-Stoke equations^24,25^. These models are often formal and theoretical, designed to be validated against physiological conditions assessed in *in vivo* experiments. For example, Oakes *et al.* employed a model based on Navier-Stoke equations to perform *in vivo* experiments on rats and determine whether lung compliance and resistance differed between physiological vs pathological conditions. Other studies have also used *in silico* lungs models to assess particle deposition and compare it with experimental findings^26^. Our model is based on the pipe-balloon model, identical to the one Maury developed for mimic *e.g.*, asthma-induced respiratory dysfunction, but it has been specifically adapted for cSCI by incorporating experimental data. In our study, the model was not constructed beforehand and then compared with experimental breathing data afterward for accuracy evaluation; instead, it was continually fitted to plethysmography experimental data. Consequently, the theoretical breathing signal is intrinsically well-aligned with the experimental data. This type of model is essential for identifying the durations of inhalation and exhalation, a critical first step for determining individualized optimal stimulation parameters.

Previous studies have proposed breathing-synchronized rNMES systems. Of note, Gollee et al. set up an algorithm to trigger abdominal stimulation (exhalation) coordinated with quiet breathing in tetraplegic patients^27^. The stimulation allowed for increased respiratory volume and capacity during exhalation, favoring cough efficiency. Another group showed similar results in terms of feasibility and reduced pulmonary complications with another system of breathing-coordinated stimulation of the expiratory muscles in patients in the acute phase of tetraplegia^28^. However, these studies mostly aim at improving abdominal muscle strength to reduce respiratory complications (including infections) and favor mechanical ventilation weaning^29^. In those with long-term mechanical ventilation, rNMES of both pectoralis and abdominal muscles might have greater impact on cough capacity and pulmonary function after cSCI. At least one study strongly suggested a good feasibility in humans, and possible functional effects^30^ in addition to direct facilitation of cough and ventilation weaning^31^. In the present study, our algorithm is not only designed for expiratory muscles but also for inspiratory ones with the ultimate goal of investigating the effect of rNMES on respiratory neuroplasticity and the potential mechanisms of action^32^. In addition, it is fully customizable and has been designed to follow the respiratory pattern, *i.e.*, spontaneous breathing, regardless of the percentage of stimulation applied during each phase or the number of rest cycles between contractions: whatever their values, the contraction and rest times will always be calculated so that the stimulation is coordinated with breathing.

The effectiveness of our algorithm relies on high quality experimental breathing signals obtained *via* plethysmography. This is critical for determining optimal parameters with a high degree of confidence. Thus, in the present study, only high-quality input signals were used (Sr of experimental data = 92%), resulting in a mean R² of 81%. These inclusion criteria inevitably led to the exclusion of some animals from the study. This illustrates a core principle of computer science: excessive noise or poor-quality input data inevitably leads to suboptimal outputs (“garbage in, garbage out”). However, translation of *Algostim* to humans may yield higher-quality data, as it is, in principle, easier to ensure that human participants remain calm during recording than it is with rodents.

More complex models such as the two-compartments model^33^ or infinite dichotomous tree model that take into consideration sub-branches and bronchial generation in the pulmonary tree have also been proposed^34,35^. These complex models would allow us to understand in-depth tracheobronchial airway and/or various properties (e.g., resistance, elastance) of the pulmonary tree accordingly^36^. In the present study, given that our primary goal was to accurately reproduce breathing signal dynamics, we opted for a simpler straightforward model to respect the parsimony principle^37,38^ and to avoid overfitting^39^. The implementation of such a conventional model to experimental data demonstrated a very good fit and required reasonable computation time. Further studies will improve the confidence of the derived optimal rNMES parameters, particularly through incorporating automatic signal screening methods to guarantee the required high-quality of the input signal^40,41^. Other computational methods related to the ODE itself (1^st^ part of *Algostim*) and/or parameter calculation procedure (2^nd^ part of *Algostim*) could also be implemented to increase the model’s fitting rate and/or parameter relevance of automatically extracted experimental data. These methods will be essential for ensuring good data reproducibility and for translating the mouse-derived algorithm to clinical studies in humans.

The current translational study has significant potential for clinical applications. Currently, only one in ten patients can be successfully implanted with a phrenic nerve stimulator to restore spontaneous breathing after SCI. The stimulation protocols based on *Algostim* replicate existing clinical practices. This non-invasive approach could benefit those with chronic cSCI, who currently have limited therapeutic options while waiting for diaphragm pacing. *Algostim* could be employed in its current form to determine its beneficial impact on respiratory neuroplasticity in rodent models of cSCI. Our acute data suggest that phase-matched stimulation may lead to improved chest excursion and greater muscle contraction compared to non-synchronized approaches. This synchronization offers a first potential advantage: it may promote greater thoracic expansion. Appropriately timed neuromuscular stimulation has been shown to induce neural plasticity and generate functional recovery in motor disorders by reinforcing natural movement patterns and strengthening residual neural circuits^42–44^. Of note, aligning peripheral stimulation with central neural activity (e.g., respiratory drive or motor intent) enhances the brain’s ability to rewire and adapt, which is directly analogous to synchronizing intercostal muscle stimulation with inspiration. Hence, while further studies are needed to confirm it, improved thoracic expansion could enhance afferent input to the central nervous system, thereby fostering neuroplastic adaptations in residual respiratory pathways.

Moreover, synchronized rNMES likely preserves spontaneous respiratory dynamics. Synchronous activation of the diaphragm and inspiratory intercostal muscles, with firing frequencies similar to those observed during spontaneous breathing, has been shown to maintain physiological respiratory patterns. By respecting the physiological timing of muscle activation, this approach allows intercostal muscles to complete their natural relaxation phase at the end of inspiration, thereby facilitating smooth exhalation^23^. This method avoids disrupting the natural respiratory rhythm, which is essential for maintaining efficient ventilation. Finally, synchronized rNMES may minimize the risk of autonomic dysreflexia (AD) that it may involuntary induce. Indeed, in SCI above T6, asynchronous or noxious stimuli can trigger uncontrolled sympathetic discharge, leading to life-threatening complications. By aligning stimulation with spontaneous breathing, synchronized rNMES could reduce the likelihood of unexpected stimuli, thereby potentially lowering the risk of AD, though this requires further validation is required^45^.

## Conclusion

*Algostim* represents an innovative computational approach combined with experimental data for synchronized stimulation of respiratory muscles, while respecting the natural respiratory patterns. With biological data now readily available and accessible, *Algostim* enables the easy delivery of personalized therapeutic interventions. Of note, in breathing disorders following cSCI, the overriding objective is to contract the inspiratory muscles during inhalation and the expiratory muscles during exhalation. Not only does *Algostim* support this challenge, but it also offers advanced customization options, including the stimulation percentage and the number of rest cycles. It has been designed to provide breathing-synchronized stimulation, regardless of the selected intensity. The reproducible and deterministic workflow facilitated by *Algostim* opens new avenues for developing therapeutic interventions that not only prevent respiratory muscles atrophy, but also facilitate coughing and improve overall breathing capacity. *Algostim* could provide a straightforward method that guarantees consensus on how the respiratory muscles should be stimulated. The current study prompts the exploration of new challenges to understand the mechanisms underlying potential respiratory neuroplasticity.

## Acknowledgments

The authors would like to thank Pr. Bertrand Maury for sharing his work on mathematical modeling of the respiratory system, Brigitte Quenet for her help with signal processing and the members of the Inserm UMR_S 1158 unit. The authors would also like to thank the SATT Lutech, Paris, ANR Recovdia, Sorbonne Université and Inserm for their support.

## Author contributions

TC: Data curation, Formal analysis, Investigation, Methodology, Software, Visualization, Writing - original draft.

NW: Validation, Writing - review & editing.

EC: Contribution on acute data: Data curation, Formal analysis, Investigation, Methodology, Writing - review & editing

AED: Methodology, Supervision, Validation, Writing - review & editing.

IV: Conceptualization, Formal analysis, Funding acquisition, Investigation, Methodology, Project administration, Resources, Supervision, Writing - original draft, Writing - review & editing.

## Data availability

Plethysmography data is available from the corresponding author upon request.

## Code availably

The source code has been granted copyright protection by the Program Protection Agency (Agence pour la protection des programmes, FRANCE) under the reference: ALGOSTIM - X22039 - SL01210 - MA00685. Please contact the corresponding author for further information.

